# Flow: a web platform and open database to analyse, store, curate and share bioinformatics data at scale

**DOI:** 10.1101/2023.08.22.544179

**Authors:** Charlotte Capitanchik, Sam Ireland, Alex Harston, Chris Cheshire, D. Marc Jones, Flora C.Y. Lee, Igor Ruiz de los Mozos, Ira A. Iosub, Klara Kuret, Rupert Faraway, Oscar G. Wilkins, Rahul Arora, Martina Hallegger, Miha Modic, Anob M. Chakrabarti, Nicholas M. Luscombe, Jernej Ule

**Affiliations:** UK DRI at King’s College London, UK; The Francis Crick Institute, UK; Goodwright Ltd.; MRC Centre for Neurodevelopmental Disorders, King’s College London, UK; National Institute of Chemistry, Ljubljana, Slovenia; Jozef Stefan International Postgraduate School, Jamova cesta 39, 1000, Ljubljana, Slovenia; UCL Queen Square Motor Neuron Disease Centre, Department of Neuromuscular Diseases, UCL Queen Square Institute of Neurology, UCL, London, UK; Yusuf Hamied Department of Chemistry, University of Cambridge, UK; Centre for Misfolding Diseases, Chemistry of Health, University of Cambridge, UK; UCL Respiratory, Division of Medicine, University College London, London, UK; Okinawa Institute of Science and Technology, Japan

**Keywords:** Bioinformatics, Nextflow, RNA-Seq, CLIP-Seq, ChIP-Seq, database

## Abstract

Ever-increasing volumes of sequencing data offer potential for large-scale meta-analyses to address significant biological questions. However, challenges such as insufficient data processing information, data quality concerns, and issues related to accessibility and curation often present obstacles. Additionally, most experimental biologists lack the time and expertise needed to independently analyse, manage and share their own data. To overcome these hurdles, we present Flow, a web-based platform that links bioinformatic analysis and database solutions with a user-friendly interface and web API. Flow currently accommodates a range of genomics methods and further DSL2-compliant Nextflow pipelines can be added via a simple JSON schema file. Deployable on local systems or cloud services, an instance is freely accessible to academic researchers at https://flow.bio.

## Background

We are in the midst of a bioinformatics data revolution, and especially large amounts of public sequencing data are available on databases such as GEO and ArrayExpress (Barrett et al. 2010; Parkinson et al. 2008). However, these databases store either raw or fully processed data that is separated from the analysis and parameters that created it, which limits our capacity for meta-analyses which require careful attention to uniform data processing and assessment of data quality. Furthermore, it is becoming increasingly difficult for any manually curated database of processed data to keep up with, not only the volume of new data being produced, but additionally the rapid developments and diversity in the experimental protocols producing data, and bioinformatic analysis tools.

Over the past decade, multiple scientific fields have faced a “reproducibility crisis” and bioinformatics is no exception, with various historical attempts at systematically reproducing analyses from published papers falling short (Baker 2016). Such a crisis is highlighted by surveys like at Biometrical Journal, where researchers identified that only six manuscripts of 56 provided both data and code (Hothorn, Held, and Friede 2009). Despite the journal requirements for data and code sharing becoming stricter, a “reproducibility iceberg” is still apparent, describing a situation in which even when data and code is provided, obstacles to reproducibility remain, such as: difficulty installing required software, different software versions, incompatibility with the operating system (OS), incomplete metadata and insufficient parameter information, among many more (Kim, Poline, and Dumas 2018).

The increased overhead in ensuring that one’s analyses are reproducible, robust and portable to different computational architectures historically presented a barrier to bioinformaticians. However, in recent years the development of bioinformatics-specific workflow languages that handle such features have lowered the costs of making computational analyses reproducible (Wratten, Wilm, and Göke 2021). These languages handle the running of parallel jobs, execute modules of code in containers which ensure the exact same computational architecture no matter where the code is run and provide an easy-to-read structure to pipelines that facilitates code sharing and team development. In this area Nextflow is particularly notable for its highly active and engaged nf-core community project: the goal is that every bioinformatics method has a “gold-standard” pipeline decided on by community convergence. The project is unique in that anyone can contribute to the pipeline development via open collaboration on GitHub and Slack (Di Tommaso et al. 2017), (P. A. Ewels et al. 2020).

Despite the greatly enhanced usability a researcher still needs to be comfortable with the command line and basic unix skills to be able to use a Nextflow pipeline. This has led to several companies releasing platforms with a graphical user interface (GUI) to give non-coders access to the latest bioinformatics workflows, this includes Nextflow Tower, latch.bio, Seven Bridges and form.bio among others, however these are mostly prohibitively expensive to the typical bioinformatics user. Additionally, Nextflow alone does not provide solutions to data management and even within the aforementioned platforms pipeline outputs are typically files in a folder, that can easily become estranged from the analysis and parameters used to generate them.

For example, within the field of crosslinking and immunoprecipitation (CLIP), which identifies protein-RNA interactions, many variations in protocols as well as a diversity of analytic approaches, have been a barrier to data integration and sharing (Hafner et al. 2021). Various CLIP databases exist, most prominently the ENCODE project containing eCLIP data (de Souza 2012) and POSTAR3, which includes the “CLIPdb” and “RBP binding sites” modules (Zhao et al. 2022). ENCODE is populated by a few specific teams in the ENCODE consortium and in terms of CLIP, is limited to eCLIP data only. POSTAR3 contains an impressive collection of CLIP datasets, but is updated manually by a single team, presenting a bottleneck to keeping the database current. It subjects different types of CLIP data to different analysis procedures, making data integration difficult. Both databases present CLIP data separately from the analysis that produced it, and without any quality control (QC) analyses, which is problematic for performing meaningful meta-analyses. With the popularity of machine learning methods, we must also be careful to ensure that training data is of high quality, to avoid the learning of technical artefacts.

These issues initially motivated us to develop a platform that would allow not only robust data analysis, but provide a database resource of ours and others’ data for general use. Here we present Flow: a web platform that enables end-to-end bioinformatics data analysis via a user-friendly GUI, by combining the power of Nextflow for reproducible analyses with modern web database solutions. Once analysed, all stages of data processing can be seamlessly shared with the community via our open database model, thus enabling the users to trace every stage of processing from raw data to final figures that can be used for manuscripts. Flow addresses existing issues in bioinformatics databases by providing code and analysis parameters alongside data objects, prioritising meaningful quality control of datasets and providing a model where anyone can contribute at any time, without having to be part of a consortium or negotiate with specific researchers, thus simultaneously relieving a bottleneck on data sharing and democratising the analysis of sequencing data for all.

## Results

### Managing bioinformatics data life cycle with Flow

Flow is a web platform designed to simplify bioinformatics data management, analysis and sharing (more details of the implementation can be found in the “Flow Architecture” section). Whilst most scientists agree that published datasets should be public and accessible, in the initial stages of data collection, analysis and refinement, many researchers would like a degree of privacy, with the option to share data and analyses with selected collaborators or labs (Ule 2020). This typically means that the initial research is conducted on personal computers and high performance computing (HPC) clusters with some sharing of code between institutes on Github and files on Dropbox or other platforms. Then, when the research is published, the data is uploaded to a public database such as NCBI GEO or ArrayExpress (Barrett et al. 2010; Parkinson et al. 2008). In contrast, Flow supports all stages of a research project, making the transition between private early work to publicly shared mature work seamless.

Different users might enter Flow at different stages of the bioinformatics data life cycle (Figure 1). For example, wet lab biologists might upload their raw sequencing data to Flow and perform analyses aimed at establishing the quality of their data, before sharing with a collaborator for more complex analyses. In this way, wet lab biologists are able to iterate their experiments until they are confident their data is high enough quality to perform meaningful deeper analysis. When the user is happy with their data it can be made publicly available as part of the open database. The extensive metadata requirements ensure that samples in the database are sufficiently annotated to allow future use by any researcher.

**Figure 1:**
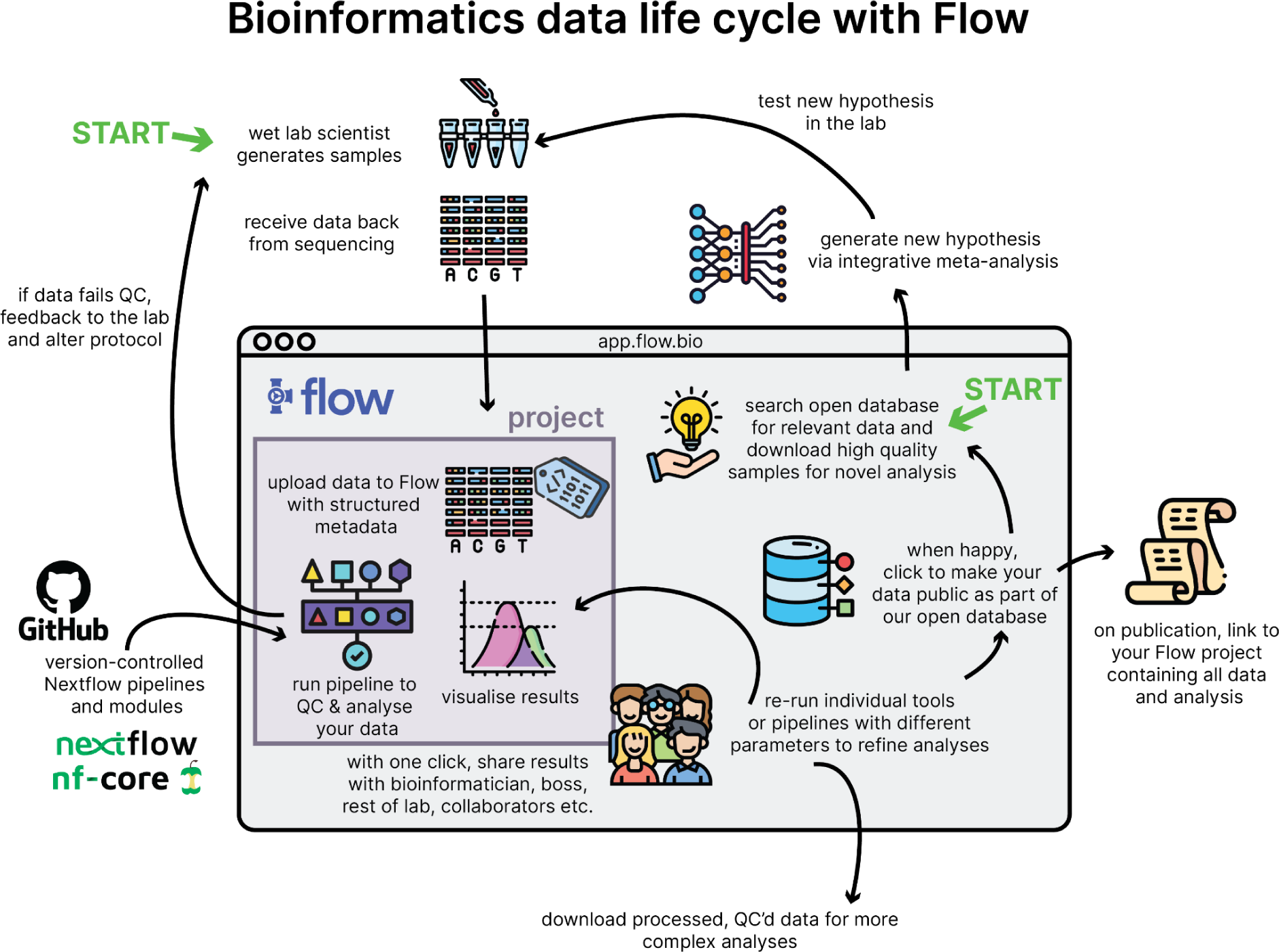
Bioinformatics data life cycle with Flow. Users can enter Flow to analyse their own samples or search for public samples relevant to their research question.

Bioinformaticians can enter Flow to search for processed data in the open database using the web platform itself, or by interacting with the database programmatically through the Flow GraphQL Application Programming Interface (API). Every sample contains detailed experimental metadata, complete logs of analysis history including versions of all software and pipelines, parameters used and runtime logs. Extensive QC information relevant to the experimental method is also available for every sample, ensuring that subsequent users of the data have all of the information needed to make informed choices about further analysis of the data.

### Data organisation and sharing within Flow

Data management on Flow is organised around several key object types, described below (Figure 2).

**Figure 2:**
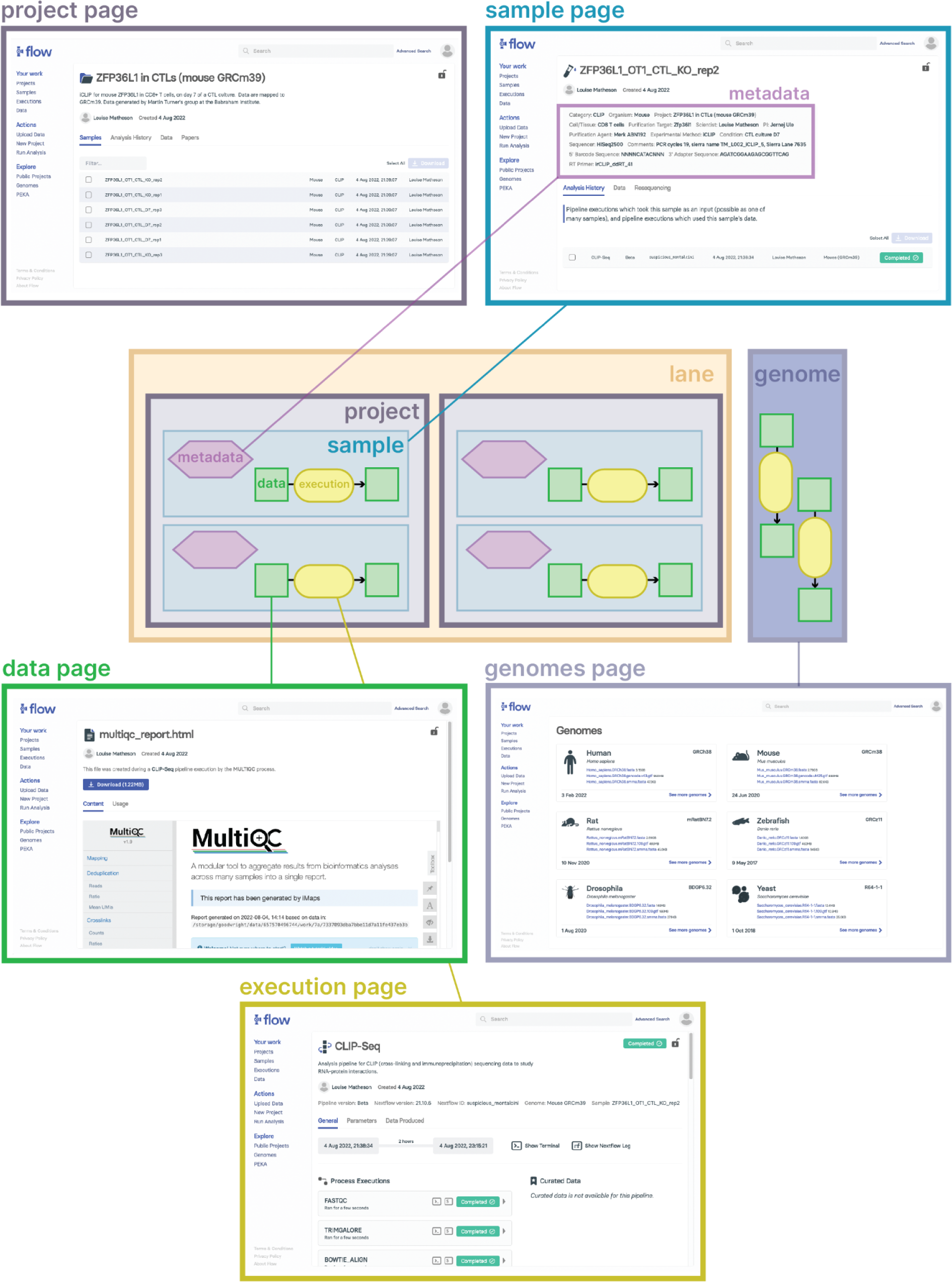
Data organisation on Flow. Data on Flow is organised based on several objects: genomes, lanes, projects, samples, data and executions. All of these objects have their own attributes which can be viewed on the web platform.

#### Projects

Projects are the top-level container for data in Flow. They are used to group related samples and other data which together contribute to a single overall research question. They may be associated with a particular published paper, and the user can decide who to share it with. Initially the user needs to decide if the project will be public - which means anyone, regardless of whether they have a Flow account will be able to see, or private - which means the project will only be accessible to the select users or groups they decide to share with.

#### Samples

In bioinformatics, typically analysis looks like a tree, where all analysis traces back to one original raw data file that came from the wet lab experiment. In Flow these initial raw data files, along with their associated metadata, are used to initialise samples. Samples can be thought of as a way of keeping all the data from a single sequencing experiment together, and easily traced back to its source. Any data generated from sample data will itself belong to that sample, so you can easily see at a glance the data belonging to a given experiment. Flow samples also contain metadata that is validated, including information about the scientist who performed the experiment and experimental conditions. Samples have a category, such as RNA-Seq, CLIP or ChIP-Seq which is used to determine which metadata is required. This is important, because metadata that is essential for CLIP data, such as the 5’ barcode sequence and targeted protein, may not be essential for other categories of data.

#### Lanes

Multiple samples may belong to the same sequencing lane, but they might not belong to the same project. Therefore, a lane on Flow will have a multiplexed FASTQ file associated with it and one or more metadata spreadsheets corresponding to samples on the lane. This makes demultiplexing easier as the user can just select a lane to pass to a pipeline, rather than manually specifying each file.

#### Executions

An execution is the record of a single run of a pipeline. It contains all the information about that run - when it was run, who ran it, what data was produced etc. Nextflow DSL2 pipeline runs consist of a series of individual processes, which run in their own self-contained environments and have their own inputs and outputs. These are chained together to form the entire pipeline execution. You can see all the processes that ran within an execution or watch them running live if the run is ongoing. Whilst the user has access to all the data that is produced, we also provide “curated data” on the execution page. Curated data represent the most commonly used outputs of the pipeline, so they can be easily accessed. Executions and data together form a network, with data nodes feeding into execution nodes, which themselves produce more data nodes. All of the benefits of running Nextflow at the command line are retained, as the terminal output and logs of the execution as a whole, and each individual process, are made available through the interface.

#### Data

In Flow, data refers to any file or folder, e.g. reads files, alignment files, log files. There are two ways to create data in Flow: data can either be uploaded by the user via the upload page, or it can be generated by executions. Some data has special labels associated with it, such as labels indicating that a file represents multiplexed reads, or is an annotation spreadsheet. Any data you generate is owned by you (whether by uploading it or by running the pipeline that generates it), with one crucial exception. If you run a pipeline which demultiplexes a multiplexed reads file into multiple demultiplexed reads files, each of those individual reads files (and data downstream of them) may have a different owner, specified in an associated annotation sheet.

Different data types are visualised on the Flow web interface to avoid unnecessary downloads. Small text files are displayed in full and previews are given of larger text files. Html files, images and PDFs are rendered inline (Figure 2: Data page), which is very useful for graphical reports generated by a pipeline, for example MultiQC reports (P. Ewels et al. 2016). All data to which a user has read-access is downloadable individually or in bulk.

#### Genomes

Analysis of bioinformatics data requires many reference files associated with genome sequences. There is a compute cost to generating these files and a storage cost, as these files can be quite large. Therefore, to prevent constantly regenerating such files, or storing many identical copies, Flow has the concept of a genome object. A genome object has a sequence fasta file at minimum, but may also contain annotations or supplementary sequence files. A genome object will have executions associated with it, for example runs of pipelines that generate the reference indexes for other pipelines. Using common genome objects also helps maintain uniformity in projects that contain many different data types and makes it easy for the user to ensure that the same annotations and sequences are used in all kinds of analysis. To enable this functionality, we have separated the Nextflow code for genome file reference generation from any Nextflow analysis pipelines that are provided on Flow. Currently six species (*Homo sapiens, Mus musculus, Rattus norvegicus, Danio rerio, Drosophila melanogaster,* and *Saccharomyces cerevisiae*) are supported on Flow, but we are working on expanding this capacity to enable support for other species.

### User accounts, groups and permissions

In order to facilitate data sharing between researchers in a safe and protected way we have established a robust system of permissions. If a user wants to perform their own analysis then they must create a free account, however public data can be viewed and downloaded even by those without an account. Beyond this individual level, users can be organised into groups. For example, a group might correspond to users from the same lab. Data, samples, executions and projects can all be shared with individual users or groups.

Every data object, execution, sample and project has a single owner - typically the user responsible for creating them. Owners have full access to the object and thus have the ability to delete them entirely. Other users can access the objects with three tiers of permissions: the lowest tier gives the ability to access the object, above this is the ability to access and edit the object, and the highest level allows users to access, edit and share the object with others (Figure 3). These can be controlled by going to the object’s page and clicking ‘Edit’ (which will only appear if you have the relevant permissions).

**Figure 3:**
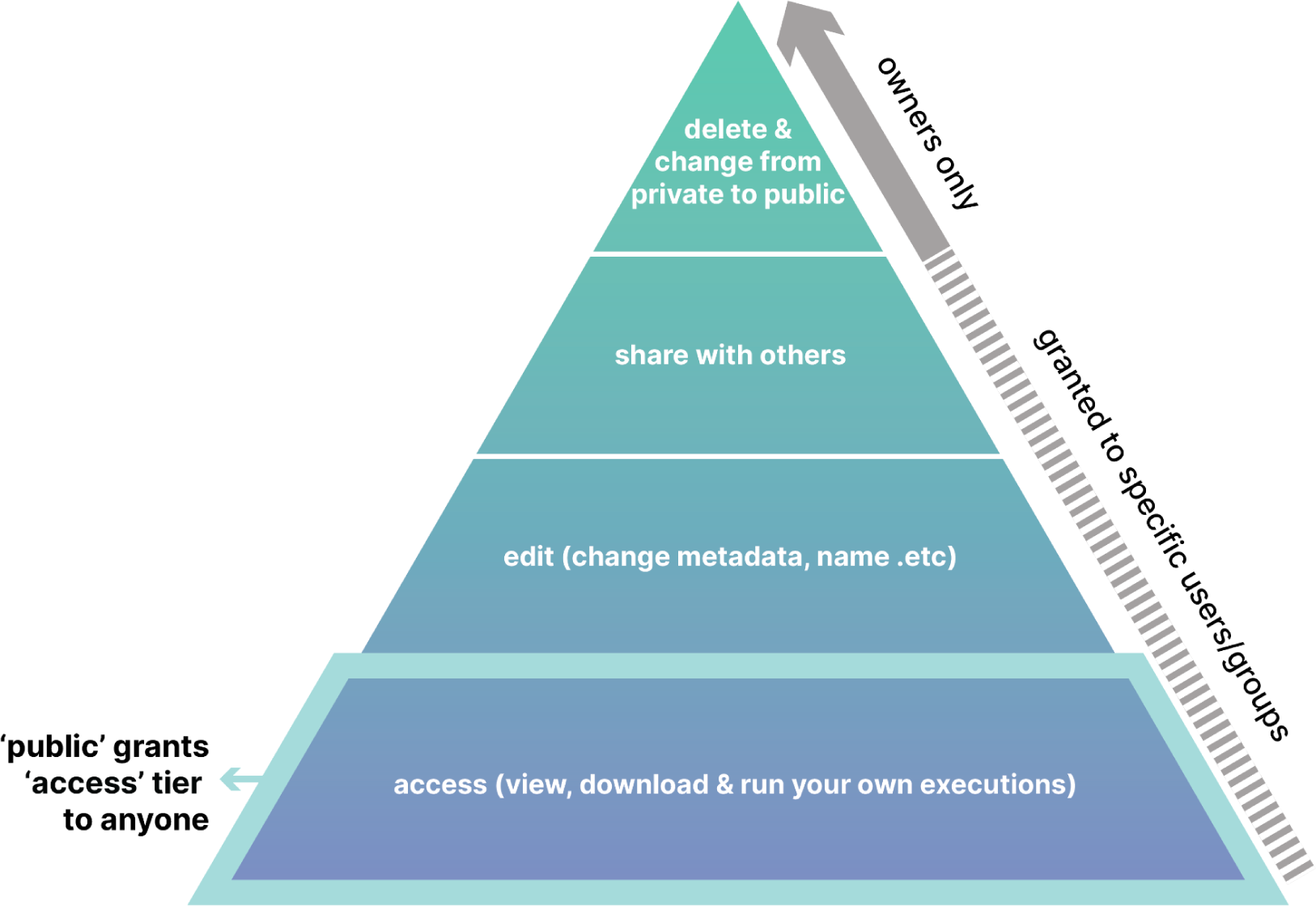
Flow permissions hierarchy. As permissions move up the pyramid, the user additionally has all the permissions of the lower tiers. Note that public objects essentially give the lowest tier of permissions to everyone, regardless of whether they have a Flow account.

All objects can also be either private (the default) or public. Public objects can always be viewed and/or used in a pipeline by anyone, regardless of whether they have been given access or even if they are signed in or not. Making an object public is equivalent to giving first tier access permissions to everyone, so even if an object is public it cannot be deleted by anyone except the owner.

In general, permissions cascade downwards - permissions given to samples will apply to any executions and data run within them and permissions given to an execution will apply to its produced data. There are two important exceptions: firstly, if the execution owner does not own some of the produced data, that data will not automatically inherit permissions from the execution. This is helpful in a situation like demultiplexing a sequencing lane, where the initialised samples downstream of the demultiplexing execution might belong to different owners, who might want to set different permissions. Secondly, if an execution is part of a sample and the owner of the execution is not the owner of the sample, then by default it will not inherit permissions from the sample (though this can easily be enabled). This is to avoid the situation in which a user may have their execution data unexpectedly made public, when the owner of the private sample they were analysing chooses to make the sample public.

### Running pipelines on Flow

Analysis on Flow typically begins with either a multiplexed or demultiplexed FASTQ, these are uploaded via the “Upload data” page. At upload time FASTQs are marked as multiplexed, demultiplexed, single-end or paired-end, which enables them to be subsequently suggested as inputs to pipelines in appropriate contexts. Available analyses can be viewed on the “Run Analysis” page and are currently categorised as “Pre-processing” pipelines, containing RNA-Seq, ChIP-Seq and CLIP-Seq pipelines, “Utilities”, containing the demultiplexing pipeline and “Modules” containing individual modules that can be run.

When running a pipeline users can choose the pipeline version and Nextflow version that should be used, to ensure reproducibility. Next, they select all the inputs to the pipeline and where these are files, appropriate suggestions are made in a drop-down menu. Users can toggle to show “advanced” options for the pipeline, enabling extensive customisation of pipeline runs, which will be recorded in the Nextflow log.

Once running, a pipeline can be monitored via its corresponding execution page, which updates in real time as processes complete. Users can additionally monitor pipeline progress by checking the provided terminal and Nextflow logs. Successfully completed processes are marked with a green icon and failed processes are marked with a red icon, alerting users to check the logging information for this process to diagnose the issue.

### An open, but orderly, database

Critical to Flow is our open database model. We could think of databases such as GEO and ArrayExpress as also being bottom-up, open databases, because anyone can contribute data to these databases, as long as it is appropriately annotated with sufficient metadata. Due to the fact they are so accessible and that journals require public data deposition, GEO and ArrayExpress don’t risk becoming “out-of-date”, users see value in uploading and sharing their own data to these repositories. The drawbacks of these databases are that they mostly only hold raw data, or final processed files, with no directly linked analysis history available or quality control information. Uploading data is often an afterthought, an additional burden at the time of publication. Further, metadata and annotation are broad enough to encompass all possible sequencing methods, meaning that the latest methodologies can sometimes be awkward to fit into this framework. In the case of CLIP data for example, it is not mandatory to include 5’ barcode information, despite the fact this is required for analysis. Other, more field-specific databases tend to be top-down, curated by a specific team, and as such become out-of-date rather rapidly. These often suffer from having little information available about the analysis history of the files or meaningful metrics describing the quality of the data, which is required for a scientist to incorporate the results into their own research.

In designing Flow, it was important for us to strike a balance between quality curation and accessibility, we wanted to make it as easy as possible to contribute data to the Flow database, whilst also ensuring that public data is described with rich enough information for it to be useful to the community. With our rigorous permissions system and project-based design, it is easy for users to make their analysis public when ready, with a simple toggle button. However, to ensure the integrity of metadata we have developed a system of metadata validation that aims to strike a balance between ensuring uniformity and being flexible enough to allow for novel experiments.

Each category of sample has its own set of metadata, with required fields that are specific to these methodologies (Table 1).

**Table 1:**
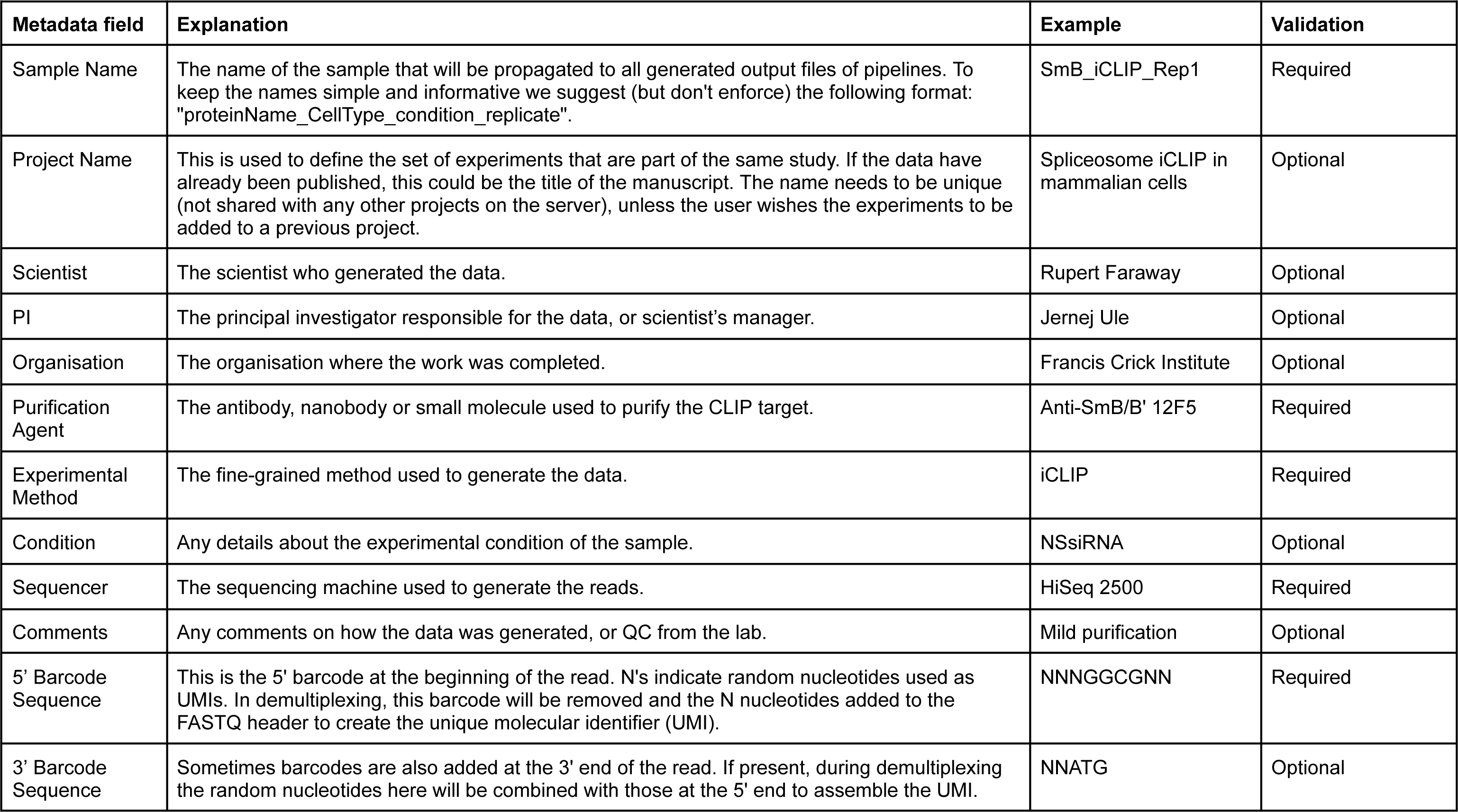
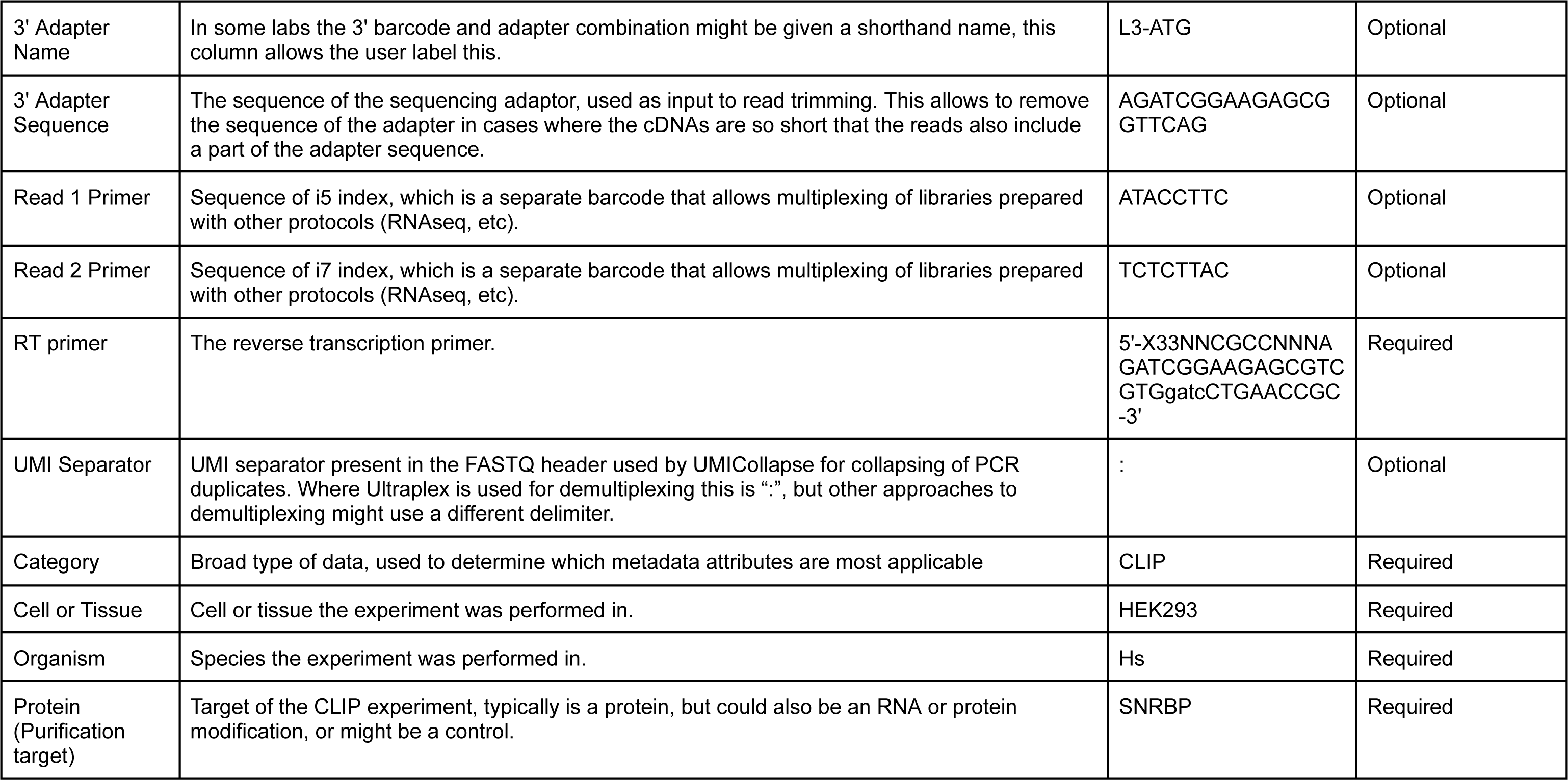
Programmatic validation criteria for CLIP sample metadata.

### Adding bespoke Nextflow pipelines to Flow is trivial with schema files

Any pipeline written in Nextflow DSL2 can be easily added to Flow with the addition of a JSON schema file (Figure 4). This schema file specifies inputs and outputs of the pipeline and any optional parameters that should be exposed on the user interface. Additionally the user can specify whether certain inputs are required or optional and whether they are considered “basic” or “advanced”, as a typical Nextflow pipeline may offer a large number of configurable parameters that are important for advanced users, but confusing to a routine user. “Advanced” inputs are made accessible on the GUI with a toggle switch.

**Figure 4:**
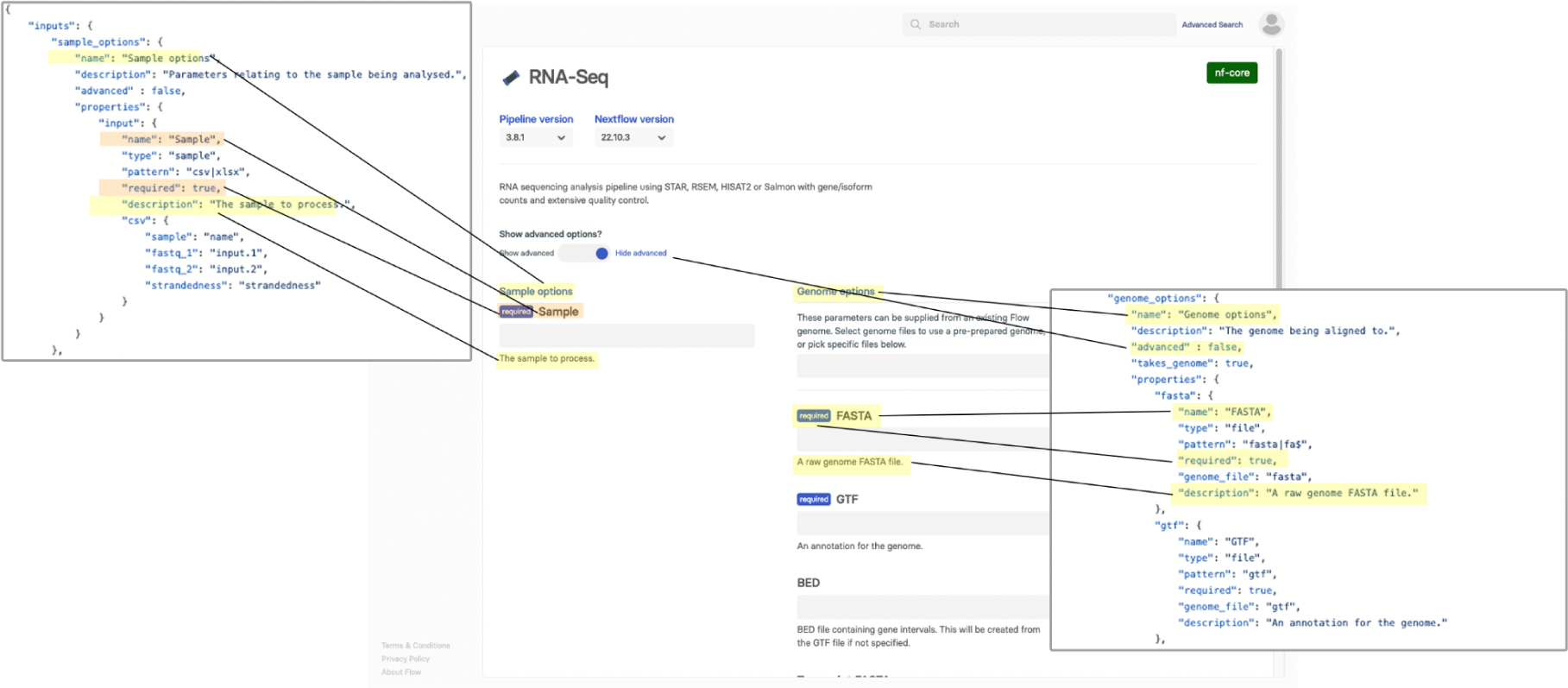
How the schema JSON file renders the user interface for a Nextflow pipeline.

Currently, addition of new pipelines to Flow requires communication with our team, primarily to ensure that robust metadata requirements are adhered to when expanding the platform to new sample types.

### Re-analysis of published STAT3 signalling data

To demonstrate the utility of Flow in managing and analysing sequencing data we created a public project containing analysis of RNA-Seq and ChIP-Seq data from a recent publication (Fike et al. 2023; https://app.flow.bio/projects/836162021371089781/; Figure 5). We were able to re-analyse the data using nf-core/chipseq and nf-core/rnaseq within the platform to get count tables for the RNA-Seq and MACS2 broad peaks for the ChIP-Seq data and crucially MultiQC reports that give quality control information such as correlation between replicate samples. The samples, analysis history, resulting final and intermediate data objects and links to the original papers are presented on the project page. Any researcher now has access to the full analysis history and quality control information of these datasets.

**Figure 5:**
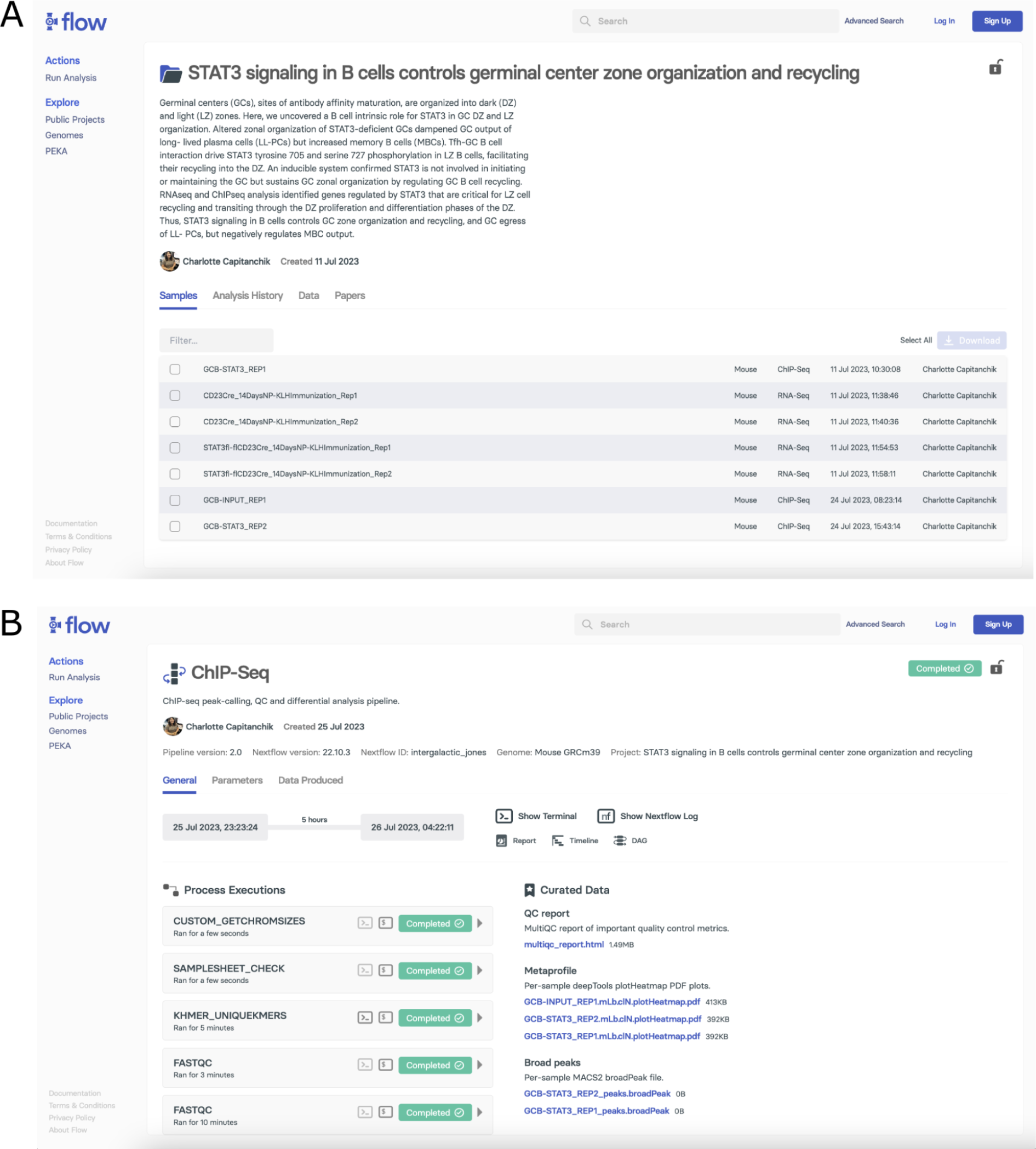
A public project showing raw data and analysis for RNA-Seq and ChIP-Seq samples from (Fike et al. 2023). A) The project page containing links to samples, analysis history, data objects and links to the papers in which the data was originally published. B) Execution page for ChIP-Seq analysis, containing all intermediate analysis steps and links to curated data such as the broad peaks, metaprofiles and MultiQC report.

## Discussion

Flow is a web-based platform we developed to address current challenges in the bioinformatics field related to reproducibility, accessibility and sharing of genomic analyses. By integrating the capabilities of the Nextflow workflow language with a modern web database and front end GUI, Flow provides an end-to-end solution for analysis and sharing of sequencing data.

We have long worked with collaborators on web servers for analysis of CLIP data (http://icount.biolab.si; http://imaps.goodwright.com), and through these efforts we increasingly realised that a linked analysis/database platform would be of broader value to many other areas of bioinformatics to enhance reproducibility and accessibility. Flow is thus designed largely to remove barriers to reproducibility of data processing, a great challenge in bioinformatics, with various historical attempts at systematically reproducing analyses from published papers falling short. Flow mitigates such problems by providing access to containerised, open-source, version-controlled Nextflow pipelines. Furthermore, by storing code and analysis parameters alongside data objects a full history of analysis is recorded, ensuring that other researchers have full access to the necessary information to reproduce an analysis.

In designing Flow we were also mindful of ensuring that the data and tools provided on the platform are findable, accessible, interoperable, and reusable (FAIR) (Wilkinson et al. 2016). To ensure findability, all data includes a unique and persistent ID, is accompanied by rich metadata and is searchable via the web platform GUI or API. Flow enhances accessibility to bioinformatics tools by providing coding-free access. Furthermore, the advanced system of user accounts, groups and permissions mean that access is tightly controlled, with the emphasis being on making data and analyses public wherever possible. Interoperability refers to how well a program or platform interfaces within the wider ecosystem. By providing access to the same Nextflow pipelines that bioinformaticians use regularly at the command line, we anticipate that Flow will fit in well with many labs’ existing practices. In the future we would like to provide integration with other resources, such as adding the capacity to import data from GEO or ArrayExpress, and export data to platforms hosting code notebooks for more complex downstream analysis. Re-usability of tools and data is paramount, to this end metadata is provided for every data object, not just per sample or project. This ensures maintenance of data provenance, which is commonly lost in bioinformatics databases. From the tooling perspective, everything is open source and tightly version controlled.

Several solutions exist for processing data using Nextflow software. Flow is complementary to these by additionally incorporating an open database model with robust metadata requirements and including features such as “curated data” to further help scientists to navigate their data. With its user-friendly GUI, Flow lowers the barriers for researchers without advanced computational skills to contribute to the data repository. The open database model allows anyone to contribute their data to the database at any time, thus democratising not only the analysis of sequencing data, but also the database contribution and curation.

Despite the advantages offered by Flow, we recognize that there are limitations to our approach. While Flow provides a user-friendly GUI, some level of bioinformatics literacy will still be necessary to correctly interpret the outputs of the analyses. Furthermore, by default, pipelines can only offer stereotyped analysis of samples. Most research projects will inevitably require a more customised, deeper bioinformatics exploration. For these reasons, Flow is not aimed at replacing bioinformaticians, but instead facilitating smoother collaborations with wet lab biologists and, much like the introduction of Nextflow itself, relieving scientists of some of the overhead required for conforming to good data practices. Furthermore, it remains to be seen how our light touch to curation will pan out in the face of large amounts of contributed data. We anticipate implementing additional features allowing researchers to curate their own and others data, based on research interest or metrics of data quality.

Flow is still undergoing rapid development and we see opportunities to expand its utility in multiple areas in the near future. Firstly, we aim to include many more pipelines and analysis options for different kinds of sequencing data in collaboration with those who are experts in these data types. Secondly, as the database grows we would like to implement analysis functionality that allows users to perform meta-analyses from within the platform, allowing users to place their own data in the context of all other data in the database from an early stage of data analysis. In both our first and second goals we see interactive visualisations as a priority, something we have already trialled on Flow with our interactive RBP motif heatmap generated from all eCLIP data using the PEKA tool (https://app.flow.bio/peka/, (Kuret et al. 2022). Finally, in the spirit of bottom-up approaches to data democracy, we are developing technology to allow Flow to be deployed in multiple locations, allowing there to be many private databases, all linked to one public-facing database. This federated approach will allow institutions to host Flow on their own HPC architecture, much more cheaply than performing analysis on the Cloud, whilst still being able to contribute to a central public data repository. By focusing our attention on how to improve and facilitate collaboration, how to present data in a biologically meaningful way and by making usability a priority, every researcher benefits.

## Methods

### Flow architecture

Conceptually, Flow operates on five hierarchical layers, where layers 3-5 have been developed entirely by us:

1. **The Bioinformatics layer:** Flow uses various established command line tools for processing bioinformatics data.
2. **The Nextflow layer:** These bioinformatics tools are organised into Nextflow pipelines, which can be run in a reproducible, containerised manner using the Nextflow software. Flow uses established nf-core pipelines (P. A. Ewels et al. 2020), with some custom ones written to nf-core conventions including demultiplexing and CLIP-Seq pipelines.
3. **The nextflow.py layer:** *nextflow.py* is a Python library for running Nextflow pipelines from Python code, and parsing the resultant log files into sensible Python objects.
4. **The API layer:** *flow-api* is a Django web app which uses *nextflow.py* internally to interact with the underlying Nextflow pipelines, and other models for organising these pipeline executions, such as collections and samples. It uses a PosgreSQL database to persist data, and exposes a GraphQL API (Ireland and Martin 2021).
5. **The web client layer:** The user will typically interact with Flow using the React frontend, which communicates with the API layer via GraphQL. There is also an official Python library, *flowbio*, which also communicates with the API via GraphQL.

The actual data analysis happens in layer 1, with each of the layers above it adding varying layers of abstraction and additional features over it. A user could analyse data at any of these layers if they so wished:

- If they wanted to use layer 1, they could just find and install the various command line programs and analyse the data directly.
- If they wanted to use layer 2, they could install Nextflow and run the specially made Nextflow workflows that encapsulate the tools in layer 1.
- If they wanted to use layer 3, they could install *nextflow.py* and run the Nextflow pipelines in layer 2 from a Python script, without needing to directly run Nextflow themselves.
- If they wanted to use layer 4 they could make calls to the Flow API directly, available at api.flow.bio/graphql.
- If they wanted to use layer 5, they could go to flow.bio and use the user interface, or do so programmatically using the flowbio Python package.

We anticipate that most users of Flow will utilise layer 5, interacting with Flow via the website, which makes the latest advances in bioinformatics available to researchers with limited coding ability. However, the additional layers provide functionality for advanced bioinformaticians to interact with the platform in a variety of ways programmatically.

Flow is currently hosted and deployed as a dedicated instance on the CREATE high performance computing infrastructure (OpenStack) at King’s College London with 4TB RAM and 256 CPU cores (King’s College London, 2023). Data is stored on CREATE’s Ceph storage cluster, with daily backups both on and off-site and a separate layer of backups via Amazon Web Storage (AWS).

## Declarations

## Ethics approval and consent to participate

Not applicable.

## Consent for publication

Not applicable.

## Availability of data and materials

Whilst the codebase to render the Flow web app is closed source, the most critical aspects of the app for scientists are open-source and freely available for use by the community. The Python Nextflow engine nextflow.py (https://github.com/goodwright/nextflow.py, GPL-3.0 Licence) is open source and free to use for non-commercial uses. The Flow API client flowbio (https://github.com/goodwright/flowbio, MIT Licence) is likewise open-source. Currently available Nextflow pipelines and modules are either available from nf-core or Goodwright Github repositories. Version numbers are provided on Flow so you can be certain exactly which pipeline has been run. Nf-core code includes RNA-Seq (https://github.com/nf-core/rnaseq, MIT Licence), ChIP-Seq (https://github.com/nf-core/chipseq, MIT Licence) and some individual nf-core modules (https://github.com/nf-core/modules, MIT Licence). Additionally, we have written several pipelines available on the platform, including sample demultiplexing (https://github.com/goodwright/flow-nf/tree/master/subworkflows/goodwright/demultiplex, MIT Licence) and CLIP analysis (https://github.com/goodwright/clipseq, MIT Licence).

## Authors’ contributions

CCa: Conceptualization, Methodology, Software, Validation, Data Curation, Visualization, Writing - Original Draft.

SI, AH: Methodology, Software, Resources, Validation, Writing - Review & Editing

CCh, MJ, IAI, KK, RF, OGW, AMC: Software, Validation, Writing - Review & Editing

FL, IRM, RA, MH, MM : Validation, Data Curation, Writing - Review & Editing

NML: Supervision, Funding acquisition, Writing - Review & Editing

JU: Conceptualization, Supervision, Funding acquisition, Writing - Original Draft

## Acknowledgements

We would like to thank all members of the Ule lab for their feedback and testing of Flow predecessor platforms. Additionally, we would like to thank the wider CLIP community who have also tested previous platforms and provided valuable feedback. We are grateful to the nf-core community for their work in developing high quality, open-source pipelines that we are able to support on Flow.

